# A novel method to guide biomarker combinations to optimize the sensitivity

**DOI:** 10.1101/2024.04.12.589302

**Authors:** Seyyed Mahmood Ghasem, Johannes F. Fahrmann, Samir Hanash, Kim-Anh Do, James P. Long, Ehsan Irajizad

**Author notes:** Corresponding Author James P Long, PhD, The University of Texas MD Anderson Cancer Center, 7007 Bertner Street, Houston, TX 77030, USA, Ehsan Irajizad, PhD, The University of Texas MD Anderson Cancer Center, 7007 Bertner Street, Houston, TX 77030, USA.

## Abstract

Logistic regression has demonstrated its utility in classifying binary labeled datasets through the maximum likelihood approach. However, in numerous biological and clinical contexts, the aim is often to determine coefficients that yield the highest sensitivity at the pre-specified specificity or vice versa. Therefore, the application of logistic regression is limited in such settings. To this end, we have developed an improved regression framework, SMAGS, for binary classification that, for a given specificity, finds the linear decision rule that yields the maximum sensitivity. Furthermore, we employed the method for feature selection to find the features that are satisfying the sensitivity maximization goal. We compared our method with normal logistic regression by applying it to real clinical data as well as synthetic data. In the real application data (colorectal cancer dataset), we found 14% improvement of sensitivity at 98.5% specificity.

**Availability and implementation:** Software is made available in Python (https://github.com/smahmoodghasemi/SMAGS)

## 1 Introduction

The main goal of any classification method is discriminating different groups. To evaluate the performance of an algorithm, many metrics have been developed including accuracy, sensitivity, specificity (Pierre Baldi, 2000), and the area under the Receiver Operating Curve (AUC-ROC) (Yt Van Der Schouw, 1992) (John Su, 2012). Each metric could be useful to answer a specific question and may fail to address problems embedded into the data like the imbalance of the classes. In clinical settings, dealing with binary outcomes is one of the main objectives where clinicians want to classify patients to healthy control and disease groups with a set of numerical variables and other covariates. The separation of these two groups constitutes a central objective in many biostatistical analyses (Chayakrit Krittanawong, 2020). This has led to the development of numerous statistical methods tailored to this challenge (Davide Chicco, 2020). In the context of binary classification, two metrics, sensitivity or true positive rate (TPR) and specificity or true negative rate (TNR), are among the most common measurements of the performance of a method which is focusing on either the rule in, correctly classifying the positive group, or rule out, correctly classifying negative group, concept. Sensitivity is the proportion of true positives which are correctly predicted positive. It evaluates a model’s ability to avoid false negatives and is particularly relevant when the goal is to minimize the risk of missing positive cases. A model with high specificity, and an acceptable sensitivity, indicates that the model is good at ruling out negative cases, meaning it correctly identifies most negative instances with a low rate of false positive (type II errors). In the context of diseases with low prevalence (i.e. cancers), a test with very high specificity is important because high false positive rates result in many healthy individuals undergoing unnecessary clinical procedure (Arash Jamshidi, 2022) (E. Klein, 2021) (Joshua Cohen, 2018). Prioritizing specificity is key to building trust and maximizing the benefits of these tests. Existing statistical techniques and machine learning algorithms often attempt to strike a balance between these metrics. For instance, the logistic regression goal is to find the coefficients to balance between sensitivity and specificity, by maximizing the log-likelihood function (Cyrus R. Mehta, 1995). However, we may not always be interested in achieving high accuracy (Hadjicostas, 2006) (Linnet, 1987). High value for specificity can be difficult to achieve, as logistic regression inherently seeks coefficients that mediate between these two ratios by maximum likelihood estimation.

Moreover, in (Hadjicostas, 2006), the author sought optimal cut-off points to maximize weighted sensitivity and specificity but did not calculate the optimum combination rule. The concept of partial AUC was introduced in (McClish, 1989) to accommodate various ranges of interest values for specificity. However, in (Cook, 2007) the author noted that the use of AUC might yield statistical significance but not necessarily clinical significance when assessing new markers. As a response, Osamu Komori and colleagues (Osamu Komori, 2010) employed ensemble techniques to compute partial AUC in a large dimensional space. Nevertheless, it is worth noting that none of these prior studies delved into the loss function of logistic regression as the means to obtain the answer. In this context, we have implemented a significant alteration in the loss function, tailored to the desired level of sensitivity at given the specificity values and vice versa.

Our motivation stems from the desire to determine the coefficients in logistic regression that can either optimize specificity when our objective is to rule out, or maximize sensitivity when our goal is to rule in. Additionally, our methodology may be used to perform feature selection targeted for optimizing sensitivity.

This work is organized as follows: section 2 describes the method in terms of an optimization question, in section 3 we present the results of the method when it was applied for synthetic data (section 3.1) and real data (section 3.2). We conclude with a discussion in section 4.

## 2 Methods

The main idea of Sensitivity Maximization At Given Specificity (SMAGS) is based on logistic regression. Logistic regression involves analyzing a binary outcome in relation to predictor variables. The model calculates the probability of an observation belonging to a particular outcome class. This is achieved by calculating a linear combination of the predictors using their respective coefficients. The outcome of this calculation is then transformed using the sigmoid function to determine the probability associated with each outcome class. By applying the threshold, which is the predefined specificity by default, we obtain the corresponding class label.

### 2.1 Notation and SMAGS Objective Function

Let *x*_i_ = (*x*_i1_, *x*_i2_, …, *x*_ip_)*^T^* ∈ ℝ*^p^* denote a vector of covariates and *y*_i_ ∈ {0,1} denote a binary response for observation i. There are n observations 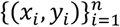. Define *y* = (*y*_1_, *y*_2_, …, *y*_n_)^T^ as the vector of response for all observations. The goal is to identify a linear separating hyperplane in ℝ*^p^* which maximizes sensitivity while obtaining at least SP specificity for the data 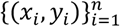. The hyperplane is defined by *β* ∈ ℝ*^p^* and *β*_0_ ∈ ℝ^1^ where an observation is predicted to belong to class 1 using the following rule:

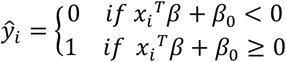

Let 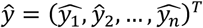 be the vector of predicted responses using the hyperplane defined by *β* and *β*_0_. Note that *ŷ* implicitly depends on *β* and *β*_0_. The sensitivity is:

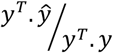

and the specificity is:

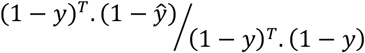

The SMAGS decision boundary (hyperplane) is:

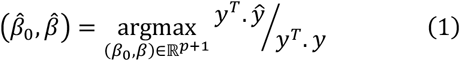

Where

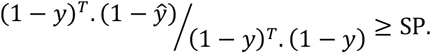

### 2.2 Optimization

In the optimization process, a diverse array of optimization techniques is employed to determine the optimal coefficients and intercept values in equation (1) that maximize sensitivity. Each optimization method yields coefficients *β* and *β*_0_., from which we compute the sensitivity. We select the set of coefficients that maximizes sensitivity for the specified specificity. The list of optimization methods are including “Nelder-Mead” (JA Nelder, 1965), “Powell” (Powell, An efficient method for finding the minimum of a function of several variables without calculating derivatives, 1964), “Conjugate Gradient” (J Nocedal, 1999), “quasi-Newton method of Broyden, Fletcher, Goldfarb, and Shanno (BFGS)” (J Nocedal, 1999), “L-BFGS-B” (R H Byrd, 1995), “TNC” (Nash, 1984), “Constrained Optimization BY Linear Approximation (COBYLA)” (Powell, Advances in Optimization and Numerical Analysis, 1994), “Sequential Least Squares Programming (SLSQP)” (Kraft, 1988) and “trust-region algorithm for constrained optimization” (J Nocedal, 1999). The first two methods are categorized as “heuristic techniques,” as they do not require the loss function to be differentiable. In contrast, the remaining methods rely on the utilization of the loss function’s derivative. Instead of computing this derivative directly, we utilize either a two-point or three-point numerical estimation approach. In instances where multiple sets of coefficients, from different optimization methods and other hyperparameter, yield the same value for the sensitivity for a given specificity (SP), our approach prioritizes the selection of the set with the lower Akaike Information Criterion (AIC) value (Akaike, 1974). This criterion facilitates the selection of a more parsimonious model, thereby enhancing the precision of our results.

### 2.3. Feature Selection Algorithm

In addition, we have formulated a feature selection algorithm rooted in the sensitivity maximization methodology outlined earlier and similar to forward stepwise selection. In this algorithm, we first fix the specificity. Then, each iteration involves the assessment of sensitivity for the present feature set augmented by a new candidate feature, encompassing all potential features. The selection of the feature that, in combination with existing features, yields the highest sensitivity is selected at each iteration. In instances where two features exhibit equal sensitivity, preference is granted to the feature associated with the lowest AIC. This iterative process continues until the desired number of features, as predetermined, is attained.

Mathematically, assume the set of original features are *F* = {*f*_1_, …, *f*_n_} and begin the feature selection method by setting specificity as SP. We compute the sensitivity corresponding to the prediction by each *f*_i_. The feature which its binary prediction, has the highest sensitivity, *f*_i_ = *v*_1_, is the first choice. In iteration *k* + 1, for a given set of features *V* = {*v*_1_, …, *v*_k_} that are already chosen, to select feature *u* from the set of remaining features {*v*_n-k+1_, …, *v*_n_}, we compute the maximum sensitivity for *V* ∪ {*u*_i_} for *n* − *k* + 1 ≤ *i* ≤ *n*. The feature *u*_j_ that yields the highest sensitivity among the augmented feature set *V* ∪ {*u*_j_} is subsequently chosen as the iteration’s selected feature.

## 3 Results

Here we present the result of our methods for two synthetic data, one with two features and one with four features, as well as real clinical data. The real data is for the Colorectal Cancer individuals, before receiving any chemotherapy. There was not also any sign of metastasis at the time of blood collection (Joshua Cohen, 2018). The synthetic data was created such that one of the features has high sensitivity and the other one has the low sensitivity, at the high value for specificity, so we can evaluate the performance of our methods at different prespecified values for specificity. For the real dataset, first we compare our method with the logistic regression with five fold cross validation and then, we perform feature selection. We compare the results at 98.5 specificity. Finally, we compare the features that our method is selected, with features selected by LASSO (Tibshirani, 1996).

### 3.1. Synthetic Data

#### 3.1.1. Two Extreme Features

In this section, we present the simulation results and conduct a performance assessment of our model in comparison to logistic regression. The synthetic dataset comprises two features, x1 and x2, each of which was generated piecewise from a uniform distribution. It follows the criteria where one of the features exhibits a high sensitivity value, while the other feature demonstrates a lower sensitivity value, both at a specificity of 98.5%. The predictive capacity of each feature concerning the binary outcome column can led to 38% sensitivity for the first feature (x1) and 3% for the second feature (x2) (Figure 1).

**Figure 1.**
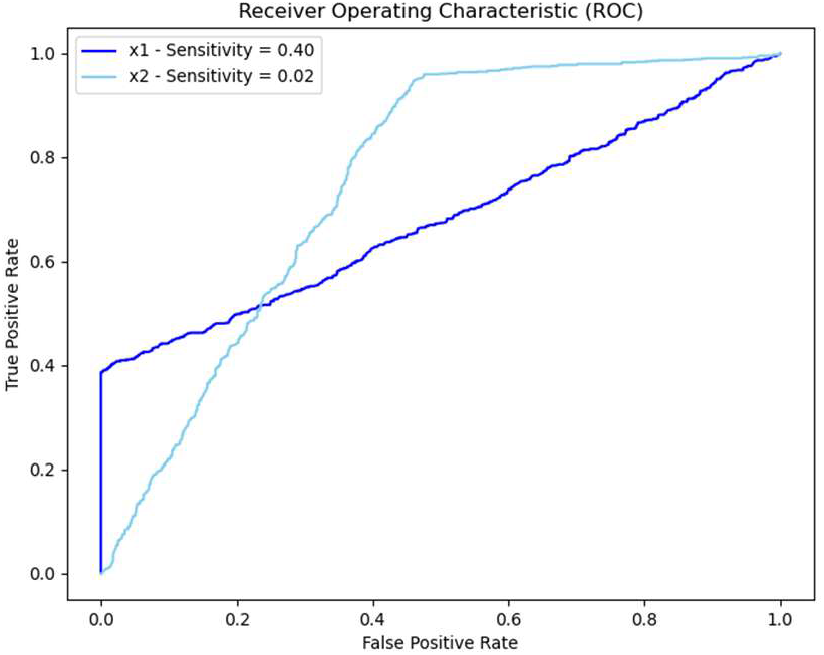
ROC of two features F1 and F2. This example shows how each feature can characterize the outcome for different value of specificity. The first one has a high value sensitivity while the other one has a low value for sensitivity.

In order to compare the two methods, we set the specificity to be equal to 98.5% in our model and find the maximum sensitivity that could be achieved. The training set includes 80% of the data with 20% of data reserved for testing. In the training set, we get 41% sensitivity from SMAGS and 33% sensitivity by logistic regression at 98.5% specificity. We should note that the goal here is maximizing the sensitivity, not the AUC. (Figure 2).

**Figure 2.**
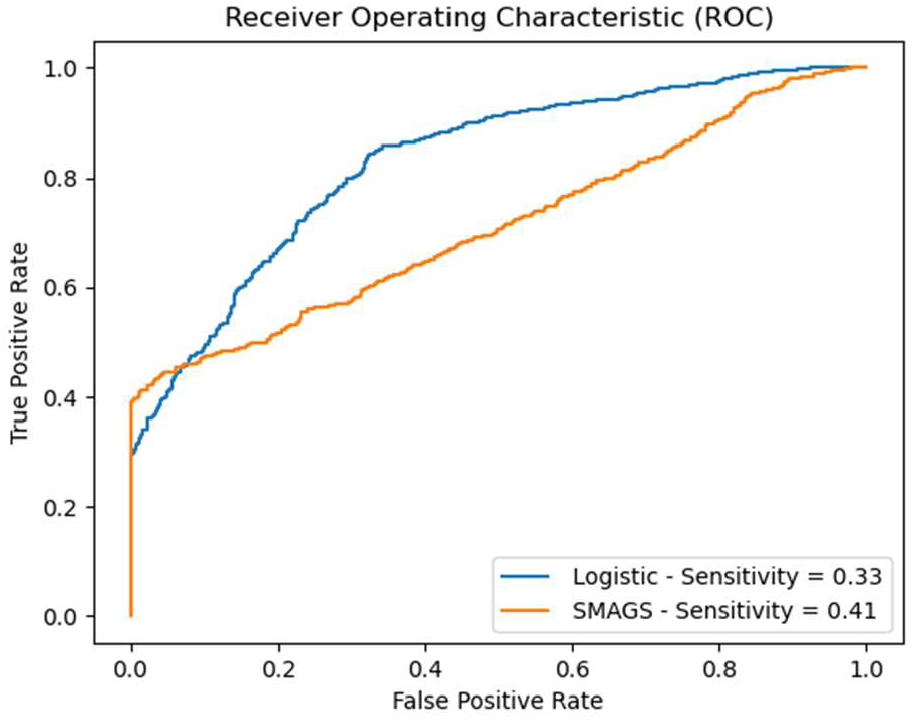
ROC of two models for training set. The value of sensitivity is computed at 98.5% specificity.

To evaluate the final performance of both models, we took the coefficients of both methods and apply it to the test set. The SMAGS has 41% sensitivity compared to 34% sensitivity of logistic regression at 98.5% specificity (Figure 3).

**Figure 3.**
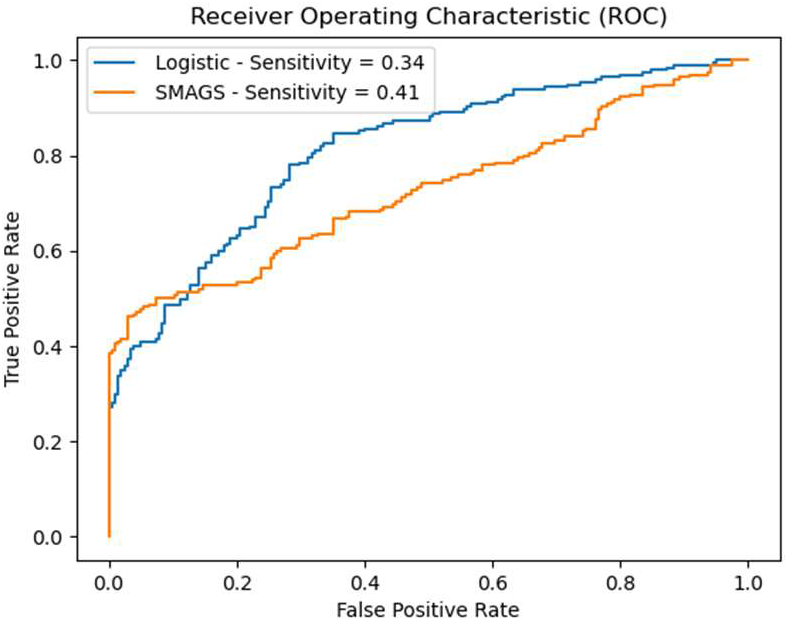
ROC of two models for test set. Comparing the sensitivity between logistic regression and SMAGS for the test set of synthetic data. The SMAGS again has higher sensitivity at the predefined, 98.5%, specificity.

Finally, we applied both methods to the entire dataset and we found 7% improvement in sensitivity at 98.5% specificity (Figure 4):

**Figure 4.**
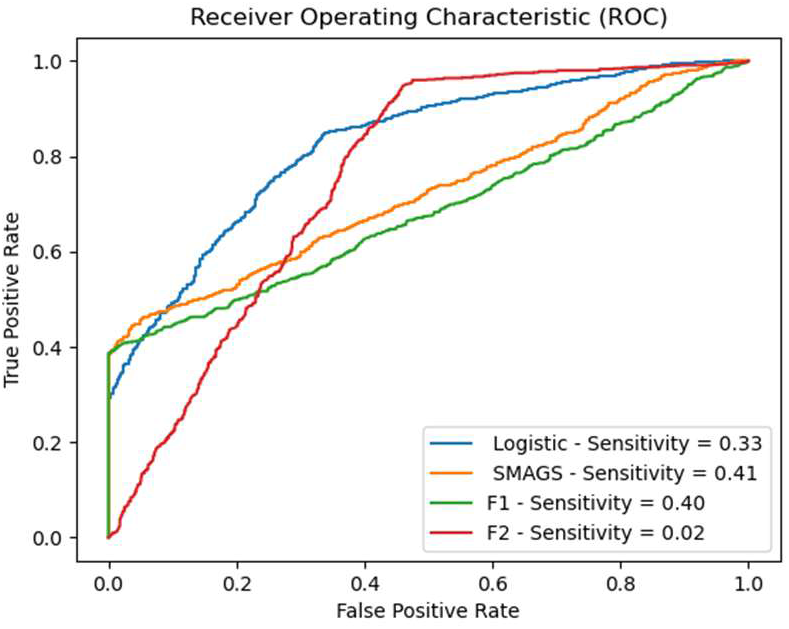
ROC of two models for test set. Comparing the sensitivity between logistic regression and SMAGS and two features for the entire set of synthetic data. The SMAGS again has higher sensitivity at the predefined, 98.5%, specificity.

#### 3.1.2. Four Features

In this section, we applied two methods to another synthetic dataset containing four features, all randomly generated from a piecewise uniform distribution. We divided the dataset into 80% for training and 20% for testing and then trained both model on the training set and apply the result for the test set. Then, we varied the specificity values from very low to very high and compared the corresponding sensitivity from logistic regression with SMAGS (Table 1) indicating a consistently higher value of sensitivity in SMAGS compared to LR at different specificity cutoffs:

**Table 1.** Comparing Sensitivity and Specificity of SMAGS and logistic regression.

| Specificity | Logistic Sensitivity (%) | SMAGS Sensitivity (%) |
| --- | --- | --- |
| 1 | 100 | 100 |
| 5 | 100 | 100 |
| 10 | 100 | 100 |
| 50 | 98.7 | 100 |
| 80 | 63.1 | 66.15 |
| 90 | 57.4 | 57.9 |
| 95 | 53.3 | 53.4 |
| 98.5 | 45.6 | 48.2 |

### 3.2. Colorectal Cancer dataset

The dataset is from published study of (Joshua Cohen, 2018) that includes 388 cases and 806 controls. There are totally of 39 proteins, where only 16 proteins have AUC > 0.6 and increased in the case group: 1-Angiopoietin-2, 2-CEA, 3-CYFRA 21-1, 4-FGF2, 5-Follistatin, 6-Galectin-3, 7-GDF15, 8-HGF, 9-IL-6, 10-IL-8, 11-Midkine, 12-Myeloperoxidase, 13-OPN, 14-Prolactin, 15-SHBG, 16-TIMP-1.

Here, we focus only on these 16 proteins. The file can be downloaded from the supplementary material of (Joshua Cohen, 2018) as well as out GitHub repository.

#### 3.2.1 Training

Similar to synthetic data, we divide the dataset into two groups, training and testing, accounting for 80% and 20% of the data, respectively. Four folds cross validation has done for SMAGS as well as logistics regression on the training part to get the coefficients. We took the average of the coefficients to assess performance on the testing set. In total, there were 908 sample in the training set and 239 samples in testing set. Out of 908 samples, 308 are cancer cases and out of 239 samples in the test set, 80 are cancer cases. For each fold in cross validation, we evaluate the sensitivity at 98.5% specificity. We see the improvement of sensitivity at 98.5% specificity for each fold, ranging from 4% (Fold 1) to 26% (Fold 4) improvement (Figure 5):

**Figure 5.**
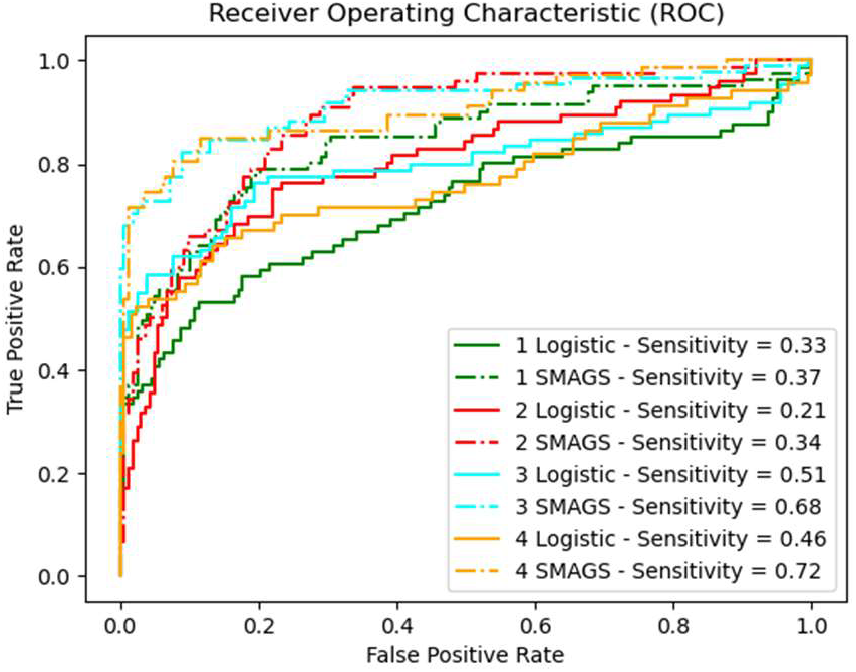
ROC of two models for each fold in training set. Comparing the sensitivity between logistic regression and SMAGS for each fold of colorectal cancer dataset. Each color represent is corresponding to one-fold, solid line is standing for logistic regression.

#### 3.2.2 Testing

Averaged coefficients from each cross validated from in the training set was applied to the testing set. We found that for the 98.5% specificity, there is 26% improvement of the sensitivity of SMAGS vs logistic regression at 98.5% specificity (Figure 6).

**Figure 6.**
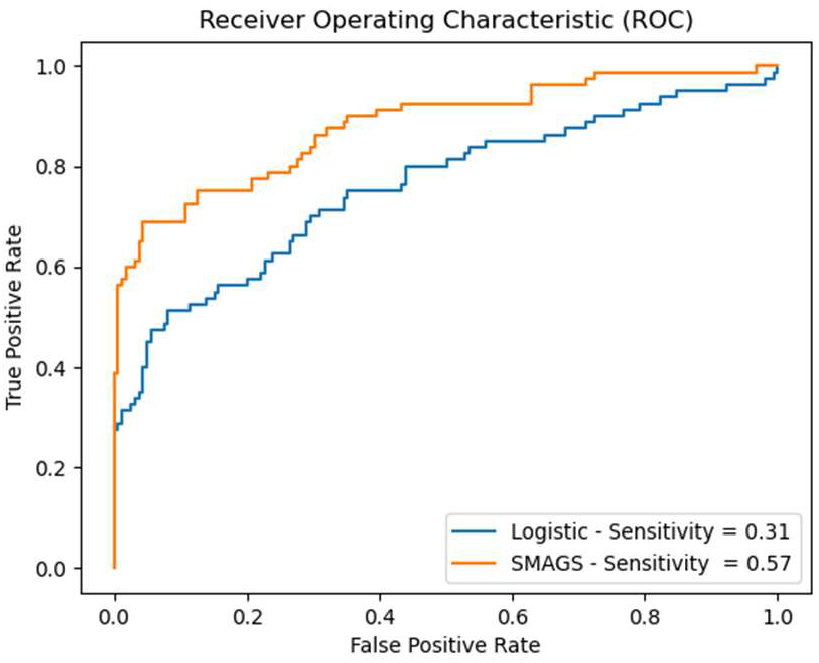
ROC of two models for test set. Comparing the sensitivity between logistic regression and SMAGS for the test set of synthetic data. The SMAGS has higher sensitivity at the predefined, 98.5%, specificity. The ROC is based on regression from the coefficient.

### 3.3. Feature Selection

In the next step, we applied the SMAGS feature selection algorithm proposed in Section 2.3 to the Cancer Seek data set. To evaluate our method for feature selection, we set the number of features to be 8 and applied it to the list of 16 proteins of colorectal CancerSeek dataset (Joshua Cohen, 2018), and following list of features was selected: 1-1-IL-6, 2-Prolactin, 3-CEA, 4-OPN, 5-Myeloperoxidase, 6-Galectin-3, 7-Angiopoietin-2, 8-FGF2. The overall performance of SMAGS vs Logistic regression when we applied the model for data with these 8 features, shows 10% improvement in sensitivity at 98.5% specificity (Figure 7).

**Figure 7.**
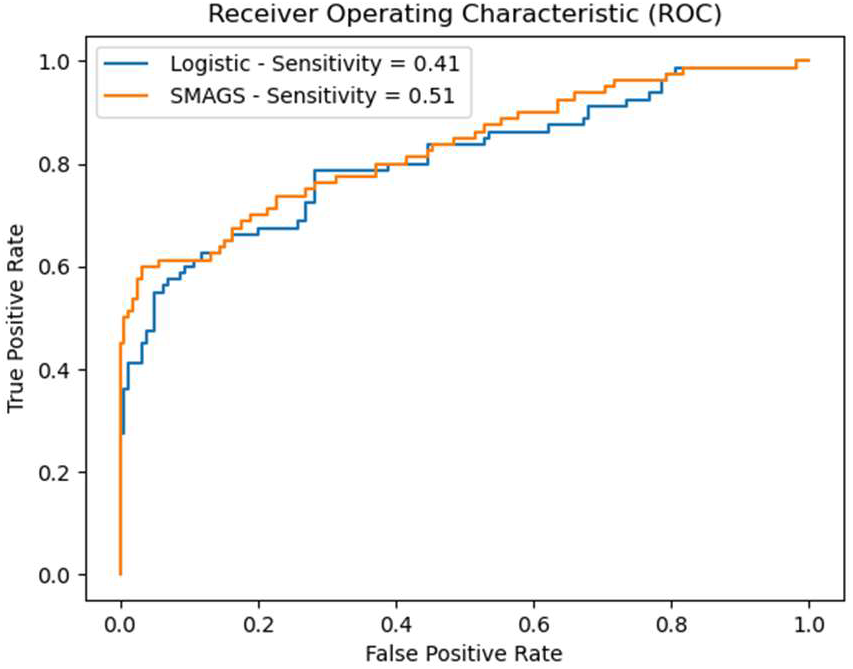
Comparison of ROC of the selected features. The features that were picked up in a feature selection process, showing 10% improvement in sensitivity at 98.5% specificity compares to logistic regression.

### 3.4. LASSO vs SMAGS

Finally, we compare the features that are selected by SMAGS with the features that are selecting with LASSO (Tibshirani, 1996) and compared their sensitivity at 98.5% specificity (scikit-learn version 1.02). We set the regularization parameter, alpha equal to 0.045, so that LASSO returns exactly 8 features with non-zero coefficients, including: 1-CEA, 2-FGF2, 3-HGF, 4-Myeloperoxidase, 5-OPN, 6-Prolactin, 7-SHBG, 8-TIMP-1. Given these 8 features, we pass them in both LASSO and SMAGS and as we expected, SMAGS has 4% higher sensitivity at 98.5 specificity (Figure 8):

**Figure 8.**
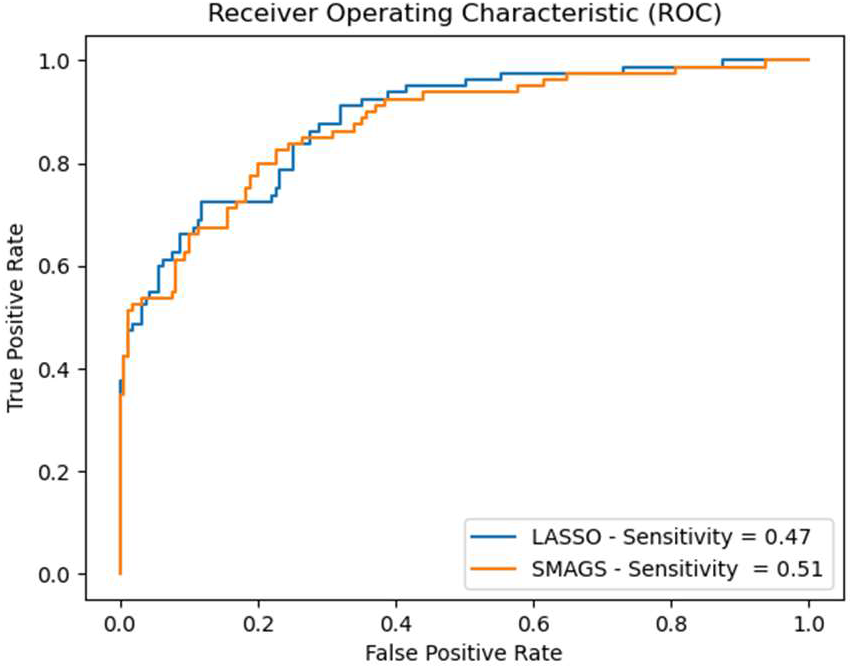
ROC of the selected features by LASSO. SMAGS also maximize sensitivity at 98.5% specificity for the eight features that are selected by LASSO. The improvement in this case is 4% at 98.5% specificity.

## 4 Discussion

We have introduced a comprehensive methodology for determining coefficients, including the intercept, applicable to regression and binary classification in clinical data. In the context of early cancer detection, our primary objective is to effectively rule-out the majority of healthy individuals while maximizing the identification of cases. Our method exhibits superior performance compared to logistic regression, demonstrated through both real-world and synthetic data.

Unlike Logistic Regression that maximizing the likelihood, our framework prioritizes the maximization of sensitivity for a pre-specified specificity. Moreover, it can selectively choose a subset of features that either enhances sensitivity (crucial for rule-in scenarios) or achieves the highest specificity (essential for rule-out situations).

As previously mentioned, we employed various optimization techniques, wherein the gradient vectors were computed using numerical methods rather than derivative functions. While numerical methods are effective in many situations, there is a possibility of failure in certain scenarios. Furthermore, this algorithm is specifically crafted to identify coefficients that maximizes sensitivity for a given specificity. This implies that if the metric of interest differs from what we have proposed here, the algorithm may not be as advantageous as other algorithms tailored to that specific metric.

Early cancer detection is a double-edged sword. While catching cancer early is vital for successful treatment, inaccurate tests can lead to downsides for patients and the healthcare system. For patients, a false positive can trigger anxiety, stress, and even unnecessary biopsies with their own risks. The healthcare system also feels the strain, with resources and attention diverted from genuine cases. The key is to strike a balance. By refining early detection methods, we can minimize these burdens and maximize the benefits of early intervention. This means saving lives and improving overall health outcomes.

## Supporting information

Colorectal Cancer Data

## 5 Supplementary Material

To access the code and the dataset that generate the graphs in this article, please visit the GitHub page at: https://github.com/smahmoodghasemi/SMAGS.

## Acknowledgements

None

## References

Akaike, H. (1974). A new look at the statistical model identification. IEEE Transactions on Automatic Control.

Arash Jamshidi, M. L. (2022). Evaluation of cell-free DNA approaches for multi-cancer early detection. Cancer Cell.

Chayakrit Krittanawong, H. U. (2020). Machine learning prediction in cardiovascular diseases: a meta-analysis. Scientific Reports.

Cook, N. (2007). Use and misuse of the receiver operating characteristic curve in risk prediction. American Heart Association.

Cyrus R. Mehta, N. R. (1995). Exact logistic regression: Theory and examples. Statistics in Medicine.

Davide Chicco, G. J. (2020). The advantages of the Matthews correlation coefficient (MCC) over F1 score and accuracy in binary classification evaluation. BMC Genomics.

E. Klein, D. R. (2021). Clinical validation of a targeted methylation-based multi-cancer early detection test using an independent validation set. Annals of Oncology.

Hadjicostas, P. (2006). Maximizing proportions of correct classifications in binary logistic regression. Journal of Applied Statistics.

J Nocedal, S. W. (1999). Numerical Optimization. Springer.

JA Nelder, R. M. (1965). A Simplex Method for Function Minimization. The Computer Journal.

John Su, J. L. (2012). Linear Combinations of Multiple Diagnostic. Journal of the American Statistical Association.

Joshua Cohen, L. L. (2018). Detection and localization of surgically resectable cancers with a multi-analyte blood test. Science.

Kostas Florios, S. S. (2008). Exact computation of max weighted score estimators. Journal of Econometrics.

Kraft, D. (1988). A software package for sequential quadratic programming. Forschungsbericht-Deutsche Forschungsund Versuchsanstalt fur Luftund Raumfahrt.

Linnet, K. (1987). Comparisson of Quantitative Diagnostic Tests: Type I Error, Power, and Sample Size. Statistics in Medicine.

McClish, D. K. (1989). Analyzing a portion of the ROC curve. Medical decision making.

Nash, S. G. (1984). Newton-Type Minimization Via the Lanczos Method. SIAM Journal of Numerical Analysis.

Osamu Komori, S. E. (2010). A boosting method for maximizing the partial area under the ROC curve. BMC informatics.

Pierre Baldi, S. B. (2000). Assessing the accuracy of prediction algorithms for classification: an overview. Bioinformatics.

Powell, M. (1964). An efficient method for finding the minimum of a function of several variables without calculating derivatives. The computer journal.

Powell, M. (1994). Advances in Optimization and Numerical Analysis. Springer.

R H Byrd, P. L. (1995). A Limited Memory Algorithm for Bound Constrained Optimization. SIAM Journal on Scientific and Statistical Computing.

Tibshirani, R. (1996). Regression Shrinkage and Selection via the lasso. Journal of the Royal Statistical Society.

Yt Van Der Schouw, A. V. (1992). ROC Curves For the Initial Assessment of New Diagnostic Tests. Family Practice, Oxford University Press.

